# Graph biased feature selection of genes is better than random for many genes

**DOI:** 10.1101/2020.01.17.910703

**Authors:** Jake Crawford, Casey S. Greene

## Abstract

Recent work suggests that gene expression dependencies can be predicted almost as well by using random networks as by using experimentally derived interaction networks. We hypothesize that this effect is highly variable across genes, as useful and robust experimental evidence exists for some genes but not others. To explore this variation, we take the *k*-core decomposition of the STRING network, and compare it to a degree-matched random model. We show that when low-degree nodes are removed, expression dependencies in the remaining genes can be predicted better by the resulting network than by the random model.

## Introduction

Many groups have proposed methods for integrating structured sources of domain knowledge with statistical models of gene expression data. Commonly used sources of domain knowledge include pathway/gene set databases and gene interaction networks. These data can be used to incorporate known interactions or correlations between gene measurements into statistical models, in order to improve prediction quality and model interpretability.

Several recent articles [1, 2, 3] have proposed conditions and metrics that a network should satisfy if used in this manner, and subsequently used these metrics to evaluate existing gene interaction databases. In [2] and [3], the authors argue that existing gene interaction networks have limited value for use as a modeling prior, relative to networks in which neighbors are chosen randomly from the genes profiled in the relevant gene expression study.

In [2], as a quality metric the authors use the AUC difference between the relevant network and a fully connected network, in which all genes are connected to all other genes (see Methods section for further details). The authors point out in Figure 3 of their paper that this value seems to be weakly correlated with the degree of the node in the network. That is, genes with highly negative AUC differences tend to have a small number of neighbors in the network, and genes with positive AUC differences tend to have more neighbors.

In this work, we sought to explore this observation in more detail.

## Methods

In [2] and [3], the authors explore the use of gene networks for feature selection in machine learning models fit to gene expression data. The authors argue that if a network is useful for feature selection on gene expression data, the relationships encoded in the network should help to predict the expression of a given gene. More specifically, for the expression of a given gene *g*_*i*_, the following should hold:

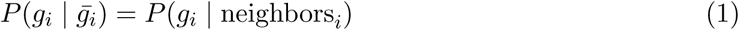

where 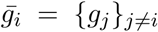 is the set containing the expression of every gene except the *i*^th^, and neighbors*i* = {*g*_*j*_ | ∃ an edge (*i, j*) in the network}. That is, if the network is informative, *g*_*i*_’s neighbors in the network are sufficient to predict its expression.

To quantify this property on real-world networks, the authors sought to predict the expression of each gene in the network based on either its neighboring genes or all other genes, using the area under the ROC curve (AUC) to compare results. Thus, the metric compared in [2] and [3] (and by us in the following analysis) is the “AUC improvement” defined as follows:

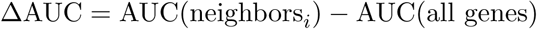

This measures the extent to which expression of the genes in the network can be predicted by their neighbors in the network, relative to a baseline prediction using all genes in the network. If the equality in (1) holds, we would expect ΔAUC to be around 0 or slightly greater, and if it does not hold we would expect ΔAUC to be negative.

For our experiments, we used the STRING network [4], which was observed in [2] and [3] to be among the best performing real-world networks. For gene expression values, we used the version of the TCGA dataset studied in [2] and [3], but all of our analyses are easily extensible to GTEx or other expression datasets.

## Results

The left panel of Figure 1 replicates Figure 3 of [2], showing that most genes with highly negative changes in AUC also had relatively low degree. Thus, we hypothesized that systematically removing genes with low degree from the network could lead to a “denoised” network that will serve as a better prior.

**Figure 1:**
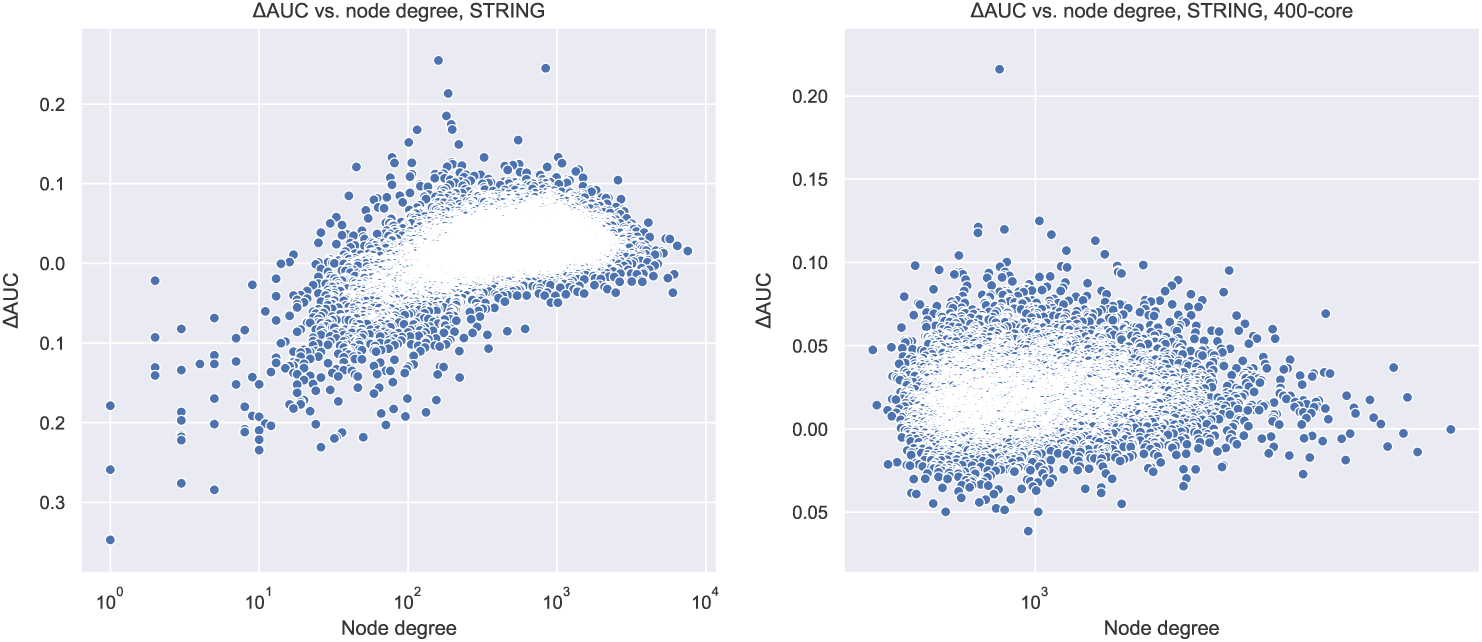
Plot of per-gene change in AUC compared with fully connected network against node degree. Full STRING network is on the left, 400-core of STRING network is on the right.

To recursively remove low-degree nodes, we took the *k*-core of the STRING network [5], for varying values of *k*. Figure 2 shows that for *k* ≈ 400 the average degree of nodes remaining in the network is maximized; thus we used the 400-core of the STRING network for further analysis. Figure 3 shows that for *k* = 400, a large part of the network is still present; approximately 3 million edges and 7200 nodes.

**Figure 2:**
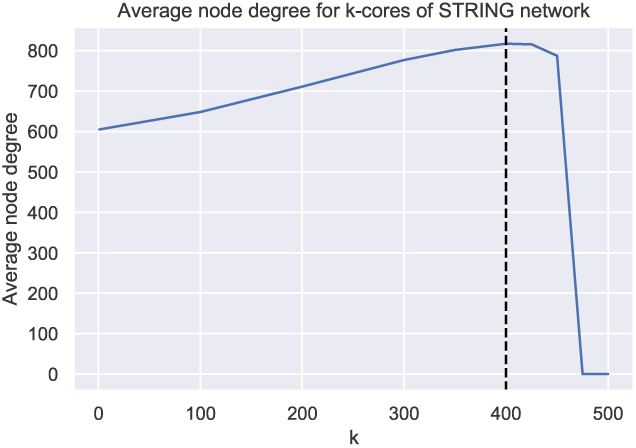
Motivating the choice of *k* = 400 for *k*-core analyses. The plot shows that for STRING, the average node degree in the network is maximized for *k* ≈ 400.

**Figure 3:**
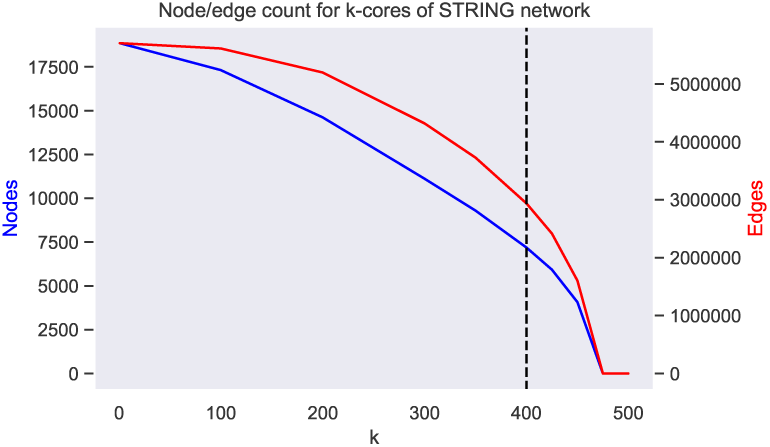
Effect of *k*-core iteration on node and edge count in the STRING network.

We then reran the experiments from [3] using the 400-core of the STRING network. In this case, as a baseline, we used a degree-matched random network as a prior (that is, for every gene left in the 400-core of the network, we randomly selected a number of other genes remaining in the network equal to the degree of the original gene, and made predictions based on the randomly chosen genes). The results are shown in Figure 4. Results comparing against R-500, R-2000, etc. graphs described in [3] are not shown here, but AUC values are similar to the degree-matched random.

**Figure 4:**
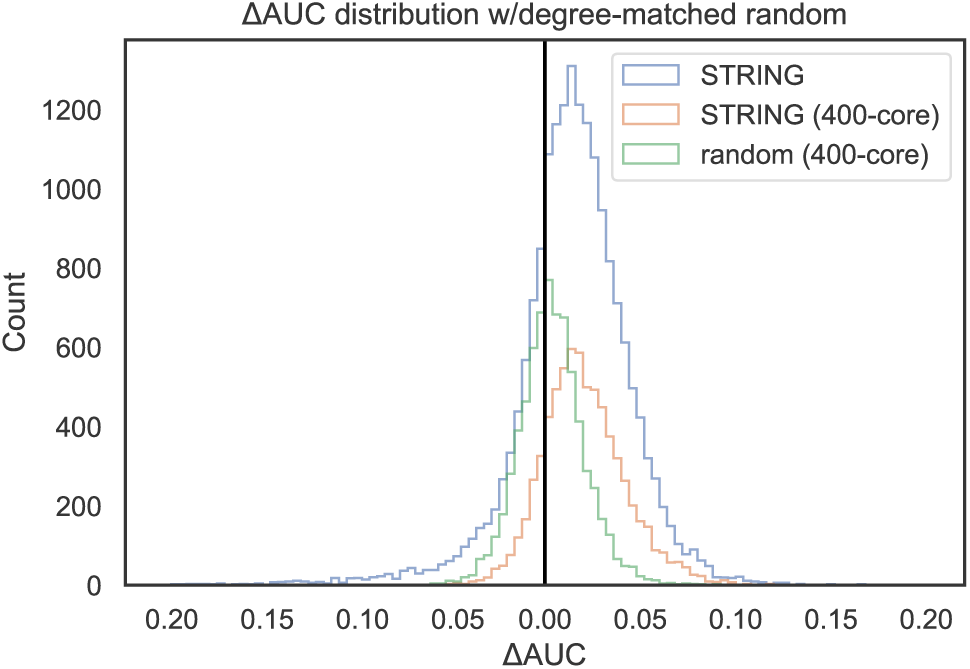
Distributions of AUC change per gene for full STRING network, 400-core of STRING network, and random network with degree matching the STRING 400-core. The more toward the right the distributions are shifted, the better the performance.

The right panel of Figure 1 shows a plot analogous to Figure 3 of [2], with the 400-core of the STRING network used instead of the full network. Taking the 400-core removes the low-degree nodes in the network, “cutting off” the poorly performing tail in the left panel.

## Conclusions

Figure 4 suggests that taking the 400-core of the STRING network does indeed appear to improve single gene inference performance somewhat, as the ΔAUC distribution is positively shifted from the degree-matched random distribution, and the “long tail” of poorly performing genes is less evident relative to the untransformed STRING network.

However, it is hard to say without further experiments whether or not this observation will be of any practical use in the context of a larger machine learning pipeline. In our *k*-core approach, weakly connected genes are removed from the network under the assumption that experimental evidence for those genes is weak and less informative; that is, we view *k*-core decomposition as a “denoising” technique. Nonetheless, it is possible that some of the genes that are removed in calculating the *k*-core simply do not have many interaction partners in truth, and predicting their expression is still biologically valuable. How these missing genes are treated in a machine learning pipeline will be of critical importance, with many possible avenues of exploration.

We chose the *k*-core approach because it is conceptually simple and has existing implementations in network processing software packages such as `networkx` and `igraph`. It is naive, however, in the sense that less connected genes are simply removed, rather than reweighted or downsampled in a more sophisticated manner. In the future, it would be interesting to try further network denoising or deconvolution approaches for this problem, such as the methods surveyed in [6] or similar. The ΔAUC metric itself also presents limitations and tradeoffs. In future work it would be interesting to compare the classification approach used here with a regression approach, in which one would directly predict expression values rather than binarizing into an above/below mean classification problem (as discussed briefly in [2]).

Our modifications to the original code and our results are available in a fork of the repository from [3], at https://github.com/jjc2718/gene-graph-conv.

